# Brain activity and connectivity attributable to nociceptive signal block

**DOI:** 10.1101/078857

**Authors:** Michael L Meier, Sonja Widmayer, Nuno M.P. de Matos, Jetmir Abazi, Dominik A Ettlin

## Abstract

Converging lines of evidence indicate that the pain experience emerges from distributed cortical nodes that share nociceptive information. While the theory of a single pain center is still not falsifiable by current neuroimaging technology, the validation of distinct brain mechanisms for acute pain and its relief is ongoing and strongly dependent on the employed experimental design. In the current study including a total of 28 subjects, a recently presented, innovative experimental approach was adopted that is able to clearly differentiate painful from non-pain perceptions without changing stimulus strength and while recording brain activity using functional magnetic resonance imaging (fMRI). Namely, we applied a repetitive and purely nociceptive stimulus to the tooth pulp with subsequent suppression of the nociceptive barrage via a regional nerve block. The study aims were 1) to replicate previous findings of acute pain demonstrating a fundamental role of the operculo-insular region and 2) to explore its functional connectivity during pain and subsequent relief. The brain activity reduction in the posterior insula (pINS) due to pain extinction was confirmed. In addition, the posterior S2 region (OP1) showed a similar activity pattern, thus confirming the relevance of the operculo-insular cortex in acute pain processing. Furthermore, the functional connectivity analysis yielded an enhanced positive coupling of the pINS with the cerebellar culmen during pain relief, whereas the OP1 demonstrated a positive coupling with the posterior midcingulate cortex during pain. The current results support the conceptual synthesis of localized specialization of pain processing with interactions across distributed neural targets.

## 1. Introduction

Ronald Melzack proposed that each bodily sensation is reflected in the human brain as a result of characteristic neural impulse patterns, and accordingly, he coined the term “neurosignature pattern for pain” (Melzack 1990). The brain regions concomitantly activated by noxious stimuli collectively have been named the “pain matrix” or “pain signature”. These include the thalamus, primary and secondary somatosensory cortices (S1 and S2), insular cortices, the anterior cingulate cortex (ACC), frontal cortices and the cerebellum (Peyron et al. 2000; Apkarian 2013; Moulton et al. 2010; Duerden and Albanese 2013). Increased activity in these areas does not necessarily imply pain selectivity but likely reflects additional unrelated processes. Regardless of whether it is nociceptive in nature, the pain matrix activity can be evoked by any salient or behaviorally relevant stimulus and might not reflect the qualitative change from non-painful to painful perception (Mouraux et al. 2011). Furthermore, accumulating evidence indicates that the pain experience is not dedicated to a specific “pain center” but rather is constructed from distributed nodes that share (anti)nociceptive information (Mano and Seymour 2015; Davis et al. 2015; Kucyi and Davis 2015). Recent advances led to a synthesis of perspectives that reconciles the localized specialization of pain processing with an appreciation for properties that emerge from interactions across distributed brain sources (Matthews and Hampshire 2016; Davis et al. 2015). Wager and colleagues convincingly demonstrated the existence of a “neural pain signature (NPS)” that discriminates between painful and non-painful brain states across many task conditions and subjects (Wager et al. 2013). However, because the NPS incorporates brain regions that are likely unrelated to nociception proper, e.g., the primary visual cortex, it raised questions about nociceptive specificity owing to a lack of evidence for nociceptive input to those brain areas (Apkarian 2013). Converging evidence from animal and human studies using central (intracortical) and peripheral stimulation suggests the existence of nociceptive afferents passing the thalamus, the cerebellum, the S1 and S2, the midcingulate cortex and the posterior insula (pINS) (Kenshalo et al. 2000; Moulton et al. 2010; Shyu et al. 2010; Garcia-Larrea 2012b; Mazzola et al. 2012a; Vierck et al. 2013; Craig 2014). In humans, the pINS and the adjacent parietal operculum (S2 region) show the uppermost preference for nociceptive signal processing, constituting a promising “core nociceptive node” (Eickhoff et al. 2006a; Garcia-Larrea 2012a; Mazzola et al. 2012b; Mazzola et al. 2012a; Segerdahl et al. 2015; Cowan 1977; Mano and Seymour 2015; Meier et al. 2015; Davis et al. 2015). Yet, while the theory of a single pain center is still not falsifiable by current neuroimaging technology, the validation of distinct brain mechanisms for acute pain and its relief is ongoing and highly dependent on the employed experimental design. Various confounding and pain-unrelated effects, such as the magnitude estimation associated with the cognitive and/or motor aspects of pain intensity rating, might blur the pain-associated neural effects (Baliki et al. 2009). Additionally, most pain studies performed categorical comparisons among different stimulus strengths, covering noxious and non-noxious stimulus ranges (Coghill et al. 1999; Brugger et al. 2012; Meier et al. 2012; Wager et al. 2013). Therefore, the neural substrate of such comparisons might be influenced by the diversity of stimulus strengths. Although efforts have been made to control for such effects by means of advanced statistical modeling (Oertel et al. 2012), it is challenging to design a proper experimental paradigm with the aim of enhancing the pain-related attribution of neuroimaging findings. Recently, we opted for an alternative approach to clearly differentiate painful from non-pain perceptions without changing stimulus strength (Meier et al. 2015). In this approach, the noxious stimulus intensity applied to a tooth was kept constant whereas the nociceptive signaling was interrupted by a local analgesic during functional magnetic resonance imaging (fMRI). In support of the current pain neuroimaging literature, a significant and exclusive reduction in brain activity in the pINS, after blocking the nociceptive barrage, was observed. The current report is based on an identical approach, but it additionally includes a placebo condition to replicate or dismiss the presumed pain processing preference of the operculo-insular cortex. Furthermore, its potential neural interactions with other brain regions were assessed during pain and its relief using generalized psychophysical interactions (gPPI). Herein, we expected to find novel and distinct functional connectivity measures that are strongly related to pain and its relief.

## 2. Methods

### 2.1 Subjects

In addition to the 14 subjects described in our previous report (Meier et al. 2015), the current report involves data from a total of 28 subjects (mean age = 27.32, SD = 7.43). The right-handed male subjects were recruited by an advertisement published on an online marketplace (www.marktplatz.uzh.ch) and were enrolled after having given informed written consent. The subjects were randomly assigned (using the random function implemented in Microsoft Excel) to two groups: one group (Group A) received a dental anesthetic (see 2.2.3), whereas the other group (Group P) served as a control by receiving a placebo (NaCl). The groups were age-matched (independent two-Sample t-test, t = 1.36, p = 0.18). The subjects received 50 Swiss francs per hour. The study was conducted according to the Declaration of Helsinki and was approved by the Ethics Committee Zurich, Switzerland.

The exclusion criteria were systemic disease, a history of allergy to the components of the local anesthetic solutions, local anesthesia at least 2 weeks before the experiment, caries, large restorations, periodontal disease, dental anxiety or a history of trauma or sensitivity of the mandibular canines. Dental anxiety was assessed by the Dental Anxiety Scale (DAS) questionnaire (Corah 1969). The mean DAS score was 6.57 (score range [4–11], SD = 1.95), indicating no dental anxiety in any subject, and did not differ between the groups (independent two-sample t-test, t = 0.08, p = 0.93). Alcohol was prohibited for 12 h before the experiment. The fMRI measurements were performed between 1 p.m. and 9 p.m.

### 2.2 Materials

#### 2.2.1 Dental splint

The mandibular splints were constructed from impressions made of Blu-Mousse (Blu-Mousse is a fast-setting vinyl polysiloxane material produced by Parkell, Inc., 300 Executive Drive, Edgewood, NY 11717, USA). Stainless steel electrodes were embedded in each splint at the labial and palatal centers (they served as anode and cathode) of the left and right mandibular canine. A small portion of a specifically prepared contact hydrogel was placed on the anode and cathode to minimize electrical resistance during stimulation. Furthermore, particular care was taken that the splints did not evoke pain or discomfort (Meier et al. 2015).

#### 2.2.2 Electrical stimulation

The “Compex Motion” system has been proven to evoke reliable sharp and pricking pain sensations and is described in details elsewhere (Keller et al. 2002; Brugger et al. 2012; Brugger et al. 2011; Meier et al. 2014; Meier et al. 2015). The Presentation^®^ software (http://www.neurobs.com/presentation) was used to control the experimental protocol. Shielded wires were used to avoid the radiofrequency contamination by the stimulation current.

#### 2.2.3 Dental anesthetic and placebo

For the anesthetic mental nerve block, articaine was used, which is currently the most common local dental anesthetic in Europe and has a long history of success (Cowan 1977). Articaine (4-methyl-3-[2-(propylamino)-propionamido]-2-thiophene-carboxylic acid, methyl ester hydrochloride) blocks nociceptive input by binding reversibly to sodium channels and subsequently reducing sodium influx (Becker and Reed 2012). The pulp analgesia lasts for one to two hours. Since small trigeminal fibers are generally more susceptible to local anesthetic solutions than thickly myelinated fibers, differential sensitivities are commonly observed in clinical dentistry as patients may remain disturbed by a sense of pressure despite complete analgesia (Becker and Reed, 2012). The pain offset times reported in the current study are in line with other studies reporting pulpal anesthesia onsets and the related inter-subject variability (Chumbley and Friston 2009; Kambalimath et al. 2013). In the current study, 0.6 ml of 4% solution containing 1:100,000 epinephrine was injected at the left mental foramen by a single trained and blinded dentist (JA) according to the technique described by Schwenzer and Ehrenfeld (Schwenzer and Ehrenfeld 2009). For the placebo, an equal amount of 0.9% saline solution (NaCl) was used.

### 2.3 Psychophysical assessments

#### 2.3.1 Outside MR scanner

Within 3-6 weeks prior to the fMRI experiment, the noxious intensity (NI) was determined and the dental anesthetic was injected. Subsequent stimulations were performed in separate sessions where subject reports regarding stimulus perception were recorded. Accompanied by detailed instructions, this procedure allowed for familiarizing the subjects with the anesthetic injection to minimize any arousal/anxiety effects and to guarantee a timely synchronized procedure across all subjects during the fMRI measurements.

#### 2.3.2 Inside MR scanner

While the subjects were positioned in the MR scanner, the sensory detection threshold (SDT), pain detection threshold (PDT) and NI were individually determined by applying an ascending method of limits. The left lower canine was stimulated with increasing intensity (1-mA steps) at randomized intervals between 8 and 12 s with a duration of 1 ms. The subjects were asked to indicate the SDT and PDT by pressing the alarm bell of the MR system. The NI was determined by further increasing the stimulus strength until the subject rated a “5” (corresponding to a painful, but tolerable perception) on a verbally instructed 11-point numeric rating scale (NRS) by pressing the alarm bell. The instructed NRS endpoints were “no pain” and “worst pain imaginable.” All threshold determinations (SDT, PDT and NI) were repeated three times, and the mean NI was applied in the subsequent stimulation paradigm. Furthermore, the pain quality was assessed by the verbal descriptors of “pricking,” “dull” and “pressing.” These three descriptors have demonstrated discriminative properties to distinguish between A-delta- and C-fiber-mediated pain (Beissner et al. 2010).

In a first phase (phase 1), the lower left canine was stimulated 30 times with the predefined NI at randomized intervals of 8 to 12 s. Afterwards, the subjects were instructed that they would receive either a dental anesthetic or a saline solution with no anesthetic effects (placebo). For the purpose of the injection, the original MR scanner bed position was memorized by the MR system and subsequently modified for an optimal local anesthesia setting followed by a submucosal injection (Figure 1) of either 4% articaine (Group A) or 0.9% NaCl (Group P). At the same time, the duration of the injection, including the exact re-positioning of the scanner bed, was maintained below one minute.

**Figure 1.**
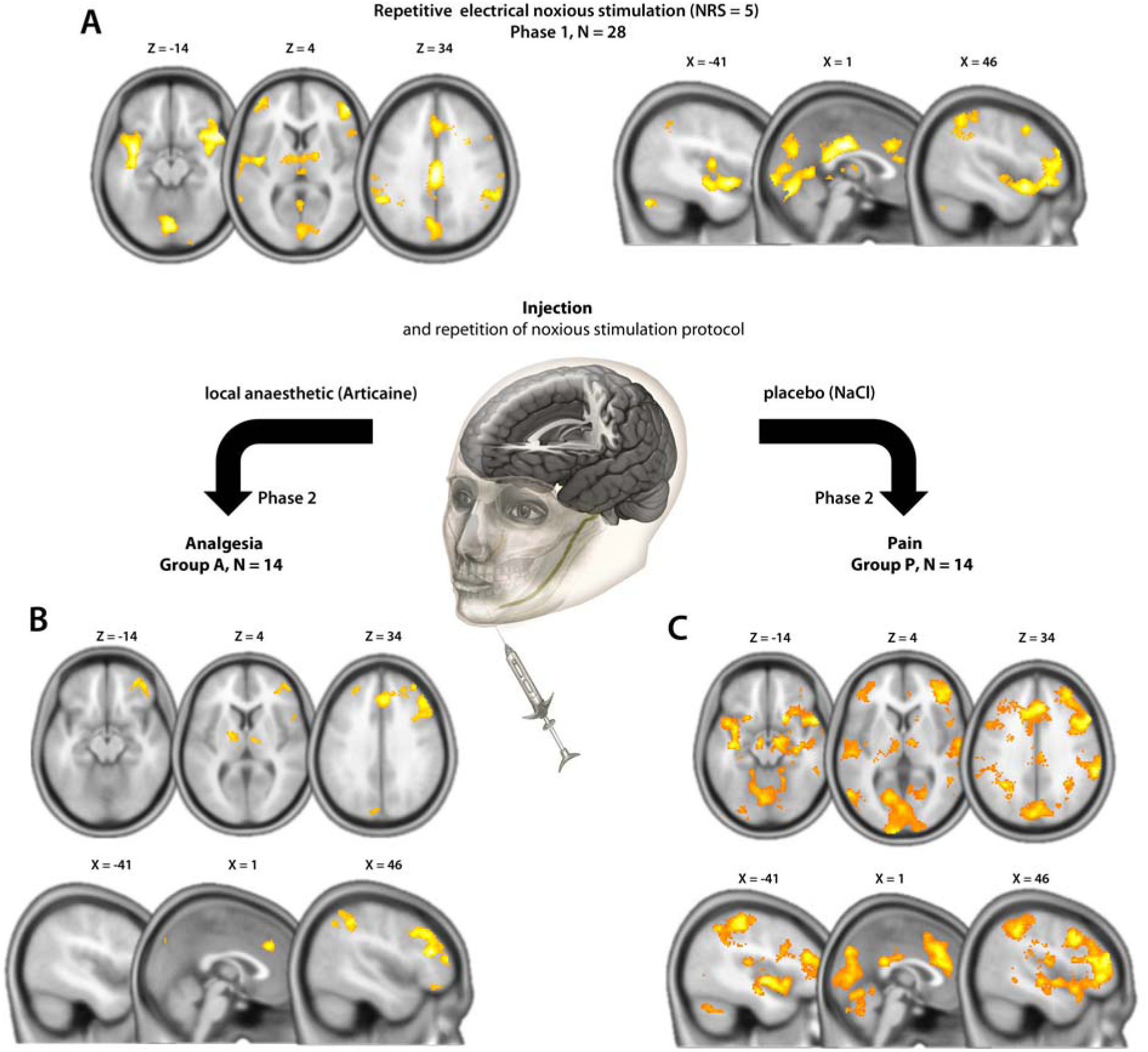
A. Overall brain activity results during the repetitive noxious stimulation in phase 1 (N = 28, one sample t-test). B. Post-Injection (articaine, Group A, N = 14) brain activity during pain relief. C. Post-injection brain activity (NaCl, Group P, N = 14). All results are thresholded with q(FDR)<0.05, cluster-based corrected.

Immediately following the injection of articaine or placebo, the subjects continued to receive repetitive electrical stimuli with equal strength as in phase 1. In the course of this second stimulation phase (phase 2), the subjects were asked to report the possible pain offset (analgesia) by pressing the MR alarm bell once. In case of reporting pain offset, subjects subsequently received 30 stimuli with predefined NI. In the case of no further perception (complete anesthesia), the subjects were asked to press the alarm bell twice.

### 2.4 Image acquisition

Functional and anatomical scans were obtained using a 3-T Phillips Ingenia scanner with a 15-channel receive-only head coil. The functional blood oxygenation level-dependent (BOLD) time series were recorded with a single-shot echo-planar imaging sequence (SENSE-sshEPI) to acquire 33 axial whole brain slices. The following acquisition parameters were used: echo time (TE) = 30 ms, repetition time (TR) = 2524 ms, FOV = 22 cm, acquisition matrix = 128 x 128, voxel size: 2.75 x 2.75 x 4.00 mm^3^, flip angle = 78° and SENSE acceleration factor R = 2.0. Using a mid-sagittal scout image, we placed 33 contiguous axial slices at 20-degree angles to the anterior-posterior commissure (AC-PC) plane, covering the whole brain. The first stimulation phase consisted of 120 functional volumes. After injection, the second stimulation phase consisted of 400 volumes in Group A and 120 volumes in Group P. A high-resolution T1-weighted anatomical image (field of view [FOV] = 22 cm, voxel size = 2.00 x 2.00 x 2.00 mm^3^, 170 slices) was recorded for each subject.

### 2.5 Preprocessing

Preprocessing of the functional brain images was conducted with the statistical parametric mapping software program SPM12 (release 6685, Wellcome Department of Imaging Neuroscience, London, UK; http://www.fil.ion.ucl.ac.uk/spm/). All volumes of the EPI sequence were corrected for slice timing, and, subsequently, a spatial realignment to the first image in the series as a reference was performed. Slices with a detected movement that exceeded 2 mm (translational) or 1° (rotational) in relation to the reference were removed. For studying the group effects, the data were normalized to the ICBM space template – European brains using seven-degree B-spline interpolation followed by smoothing with a Gaussian kernel of 8 mm full-width-at-half-maximum (FWHM) (Evans et al. 1992). While smoothing is necessary to produce reliable estimates of statistical significance using the theory of Gaussian random fields, it should be noted that the size of the smoothing kernel can influence the estimation of the true spatial extent of brain activity (Stelzer et al. 2014).

### 2.6 Statistical modeling and analysis

#### 2.6.1 Brain activity analysis

Single subject and group analyses were performed with SPM12 (release 6685). A general linear model (GLM) was applied to partition the observed neural responses into components of interest, confounders and errors (Friston 1995). An event-related analysis estimated the BOLD responses evoked by the potentially noxious stimuli by modeling them as a delta function convolved with the canonical hemodynamic function as implemented in SPM12. The stimulus duration was 1 ms with a randomized interstimulus interval between 8 and 12 s. In the 1^st^ level (single subject) analysis, all 30 noxious stimuli of phase 1 were modeled as a single regressor (“pain phase 1 regressor”) in both groups. In phase 2, all 30 noxious stimuli of Group P were identically modeled (“pain phase 2 regressor”). In Group A, the first painful stimuli and the pressing of the alarm bell were individually implemented as regressors of no interest, whereas the following 30 non-painful stimuli (“non-painful phase 2 regressor”) were modeled as regressors of interest. Additional confounders were incorporated in each analysis within the design matrix, including the six rotational and translational parameters from the rigid body transformation, obtained during the functional image realignment. Low-frequency fluctuations were removed with a high-pass filter (128 s). The serial autocorrelation of the BOLD time series was modeled using a first-order autoregressive model (AR[1]). The computed contrast maps derived from each subject were then entered into a random effects (RFX) analysis.

To properly limit the amount of false positives, whole-brain topological inference using a cluster-based false discovery rate method (FDR) based on Gaussian Random Field Theory was applied (Chumbley and Friston 2009). The cluster defining threshold was set to p < 0.001. An initial threshold of p < 0.001 is recommended to avoid the pitfalls associated with cluster-based thresholding (Woo et al. 2014; Eklund et al. 2016). Only clusters that survived the FDR correction were used for the interpretation of the results. For within-group analyses, one-sample t-tests were performed, whereas for between-group analyses, independent two-sample t-tests were used as implemented in SPM12. The variance between groups was assumed to be unequal. The error covariance components were estimated using restricted maximum likelihood (REML). The activations and deactivations associated with each regressor were tested by means of simple positive and negative t-contrasts. To uncover any possible pre-injection differences in brain activity, phase 1 was compared between groups. Furthermore, the post-injection brain activity between groups was investigated by comparing phase 2. The potential effects of post-injection temporal discrepancies due to the additional painful stimuli in Group A can be neglected because adaptive changes in the sensory experience of the electrical stimulus within the experimental time window are not present (Brugger et al. 2012; Meier et al. 2014; Meier et al. 2015). Finally, the thresholded voxel SPM t-maps were color-coded and superimposed onto the avg152T1-MNI brain using xjview (http://www.alivelearn.net/xjview8/). To determine the exact OP area of the S2 region, the SPM anatomy toolbox (version 2.2c) was used (Eickhoff et al. 2005; Eickhoff et al. 2006b).

#### 2.6.2 Functional connectivity analysis

The main advantage of the PPI analysis is that it assesses the co-variance between regions across time during a certain task, and therefore provides a test of task effects on connectivity. The generalized form of the context-dependent PPI approach (gPPI) increases the flexibility of the statistical modeling and improves single-subject model-fit, thereby increasing the sensitivity to true positive findings and a reduction in false positives (McLaren et al. 2012). Importantly, the co-variance is assessed on the neural level, which results in a change in the BOLD signal, rather than at the level of the BOLD signal, which is an indirect measure of neural activity (McLaren et al. 2012; Kim and Horwitz 2008). Therefore, a deconvolution step is mandatory to capture the neural signal on which the interaction analyses are performed. As such, the gPPI model might better capture the BOLD response and the associated underlying neural activity than a conventional task regressor. However, these analyses are not able to reveal directed links between regions (i.e., activity in region A causes activity in region B). Instead, these analyses allow inferences about the co-activation of regions across subjects and time, whereas the strength of co-activation is modulated by the task state (pain/analgesia).

For each subject, we extracted the deconvolved time course of 6 mm-spheres with peak coordinates of pain-related activity in the pINS and OP1 areas identified in the between-group brain activity analysis of phase 2 (see 3.3.2, MNI coordinates OP1: 54 -20 16, pINS: -42 -8 -4). Subsequently, separate psychological (pain/relief) and physiological regressors (time course of seed regions) and their PPI interaction, as well as the movement parameters, were included in the gPPI model. Furthermore, to avoid circularity, the main effect of the task was also modeled to detect functional connectivity effects over and above (orthogonal to) the main effect of the task (O’Reilly et al. 2012). The resulting whole-brain gPPI connectivity estimates were then evaluated using a two sample t-test implemented in SPM12. The identified clusters were considered to be significant when falling below a cluster-corrected q(FDR) < 0.05.

## 3. Results

### 3.1 Stimulus perception and pain relief

The electrical stimulus strength of the NI in phase 1 (5/11 NRS) did not differ between groups (Mann-Whitney U-test, p = 0.7). None of the subjects indicated any sensitization/habituation effects of the repetitive pain stimulus. Furthermore, all subjects reported a pricking pain perception, indicating predominantly A-delta fiber-mediated pain. Following the injection, the pain stopped at 2.8 min (SD = 3.73 min) in Group A. Although no longer painful, the stimulus was always perceived by the subjects, indicating incomplete anesthesia. In Group P, no subjects reported pain relief.

### 3.2 Brain activity results

#### 3.2.1 Overall brain activity phase 1 (pain phase)

The pooled group analysis of brain activation induced by the noxious stimulation yielded activity in several brain regions commonly reported in pain studies: insular, cingulate and somatosensory cortices, cerebellum, thalamus and frontal brain regions (q(FDR) < 0.05, Figure 1a,Table 1a).

**Table 1.**
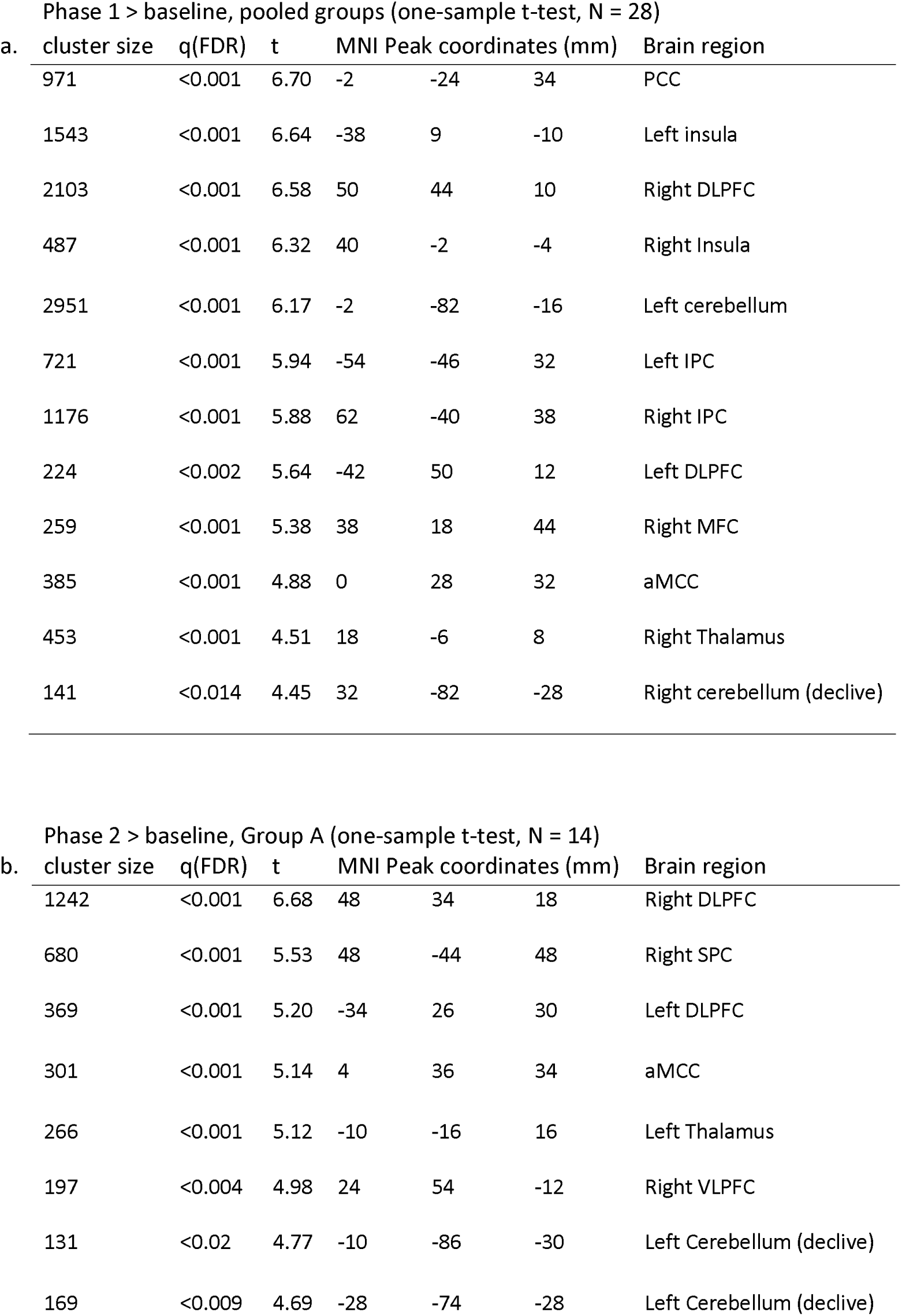
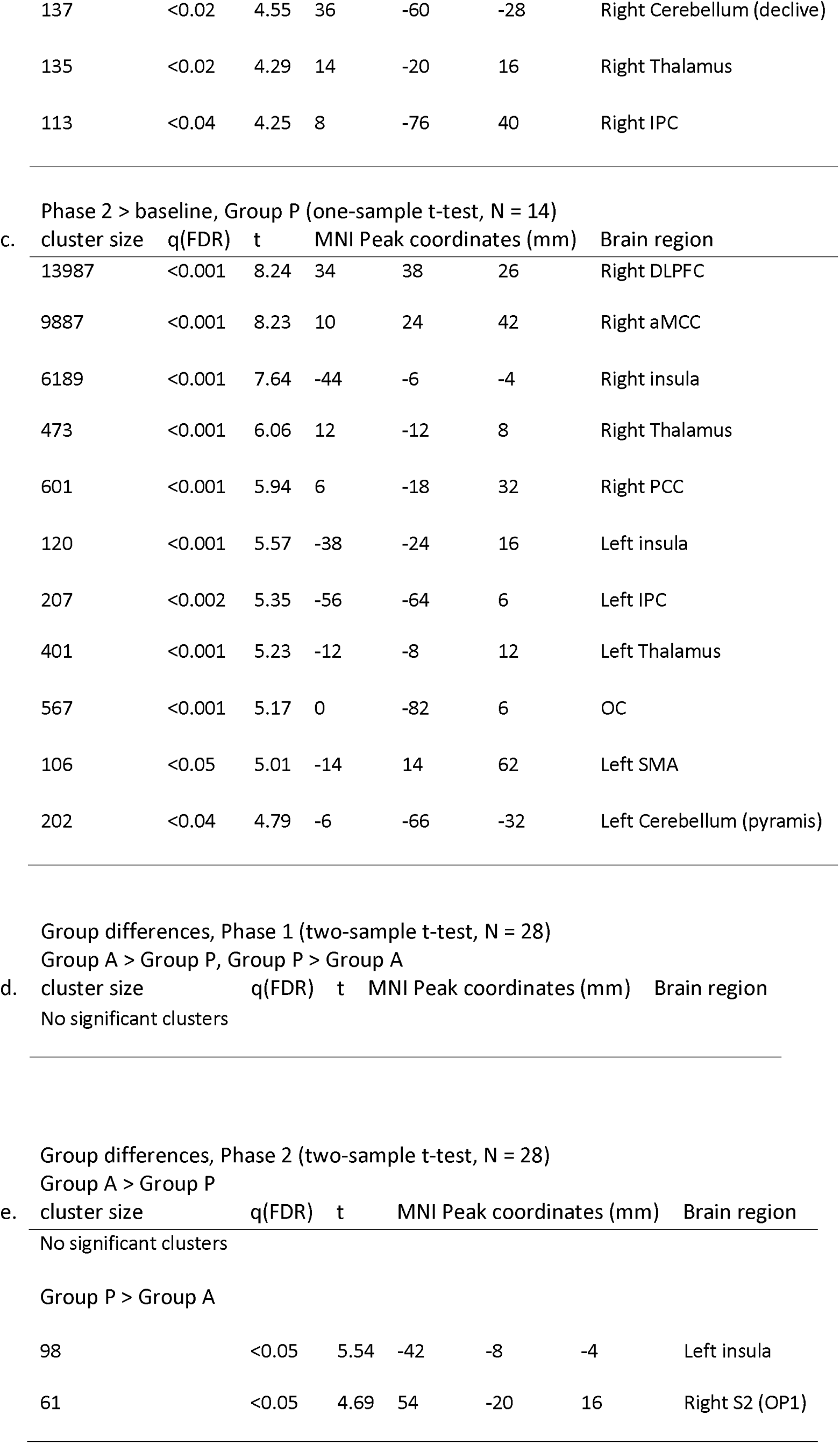
Brain activity results of Phase 1 and Phase 2. MNI = Montreal Neurological Institute, FDR = False discovery rate, PCC = posterior cingulate cortex, aMCC = anterior midcingulate cortex, DLPFC = dorsolateral prefrontal cortex, VLPFC = ventrolateral prefrontal cortex, IPC = inferior parietal cortex, SPC = superior parietal cortex, MFC = middle frontal gyrus, S2 = secondary somatosensory cortex, OC = occipital cortex

For the brain activity results of phase 2 within each group (Figures 1b and 1c) please see Tables 1b and 1c.

#### 3.2.2 Group differences

Importantly, the between-group analysis of phase 1 revealed no significant activation differences (Table 1d). In contrast, the between-group comparison of phase 2 revealed a distinct activation difference, reflected by persistent activation clusters in the ipsilateral pINS (q(FDR) < 0.05, peak MNI coordinate: -42 -8 -4, Figure 2a, Table 1e) and the contralateral S2 region, extending into the inferior parietal lobule (q(FDR) < 0.05, peak MNI 54 -20 16, Figure 2b, Table 1e) in Group P. Feeding the SPM anatomy toolbox with the peak coordinate [54 -20 16] yielded a probability of 56% (range 42-67%) for belonging to area OP1 and a 12% (range 9-13%) probability for OP4.

**Figure 2.**
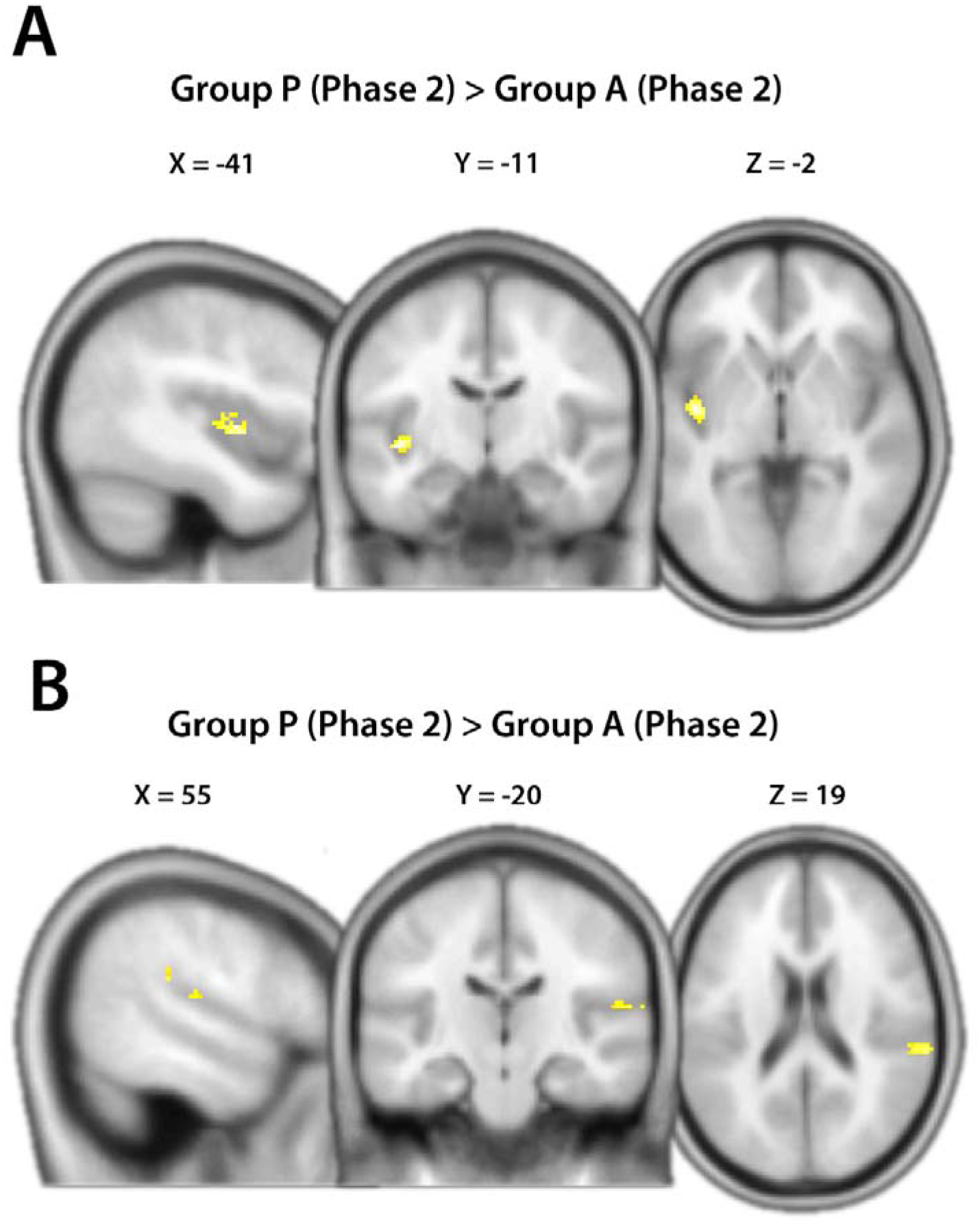
Between-group brain activity analyses (two sample t-test, N = 28). The contrast Group P (Phase 2) > Group A (Phase 2) showing brain activity strongly related to nociception revealed exclusive brain activity differences in the pINS (A) and S2 region (OP1). q(FDR)<0.05, cluster-based corrected.

### 3.3 Functional connectivity results

#### 3.3.1 Overall functional connectivity phase 1

Using the pINS as a seed (Figure 3c), the pooled group analysis yielded an enhanced positive functional connectivity during noxious stimulation to the ipsilateral insula, bilateral ventrolateral prefrontal cortex (VLPFC), S1 and S2, superior parietal cortex (SPC) and inferior frontal gyrus (IFG) (q(FDR) < 0.05, Table 2a).

**Figure 3.**
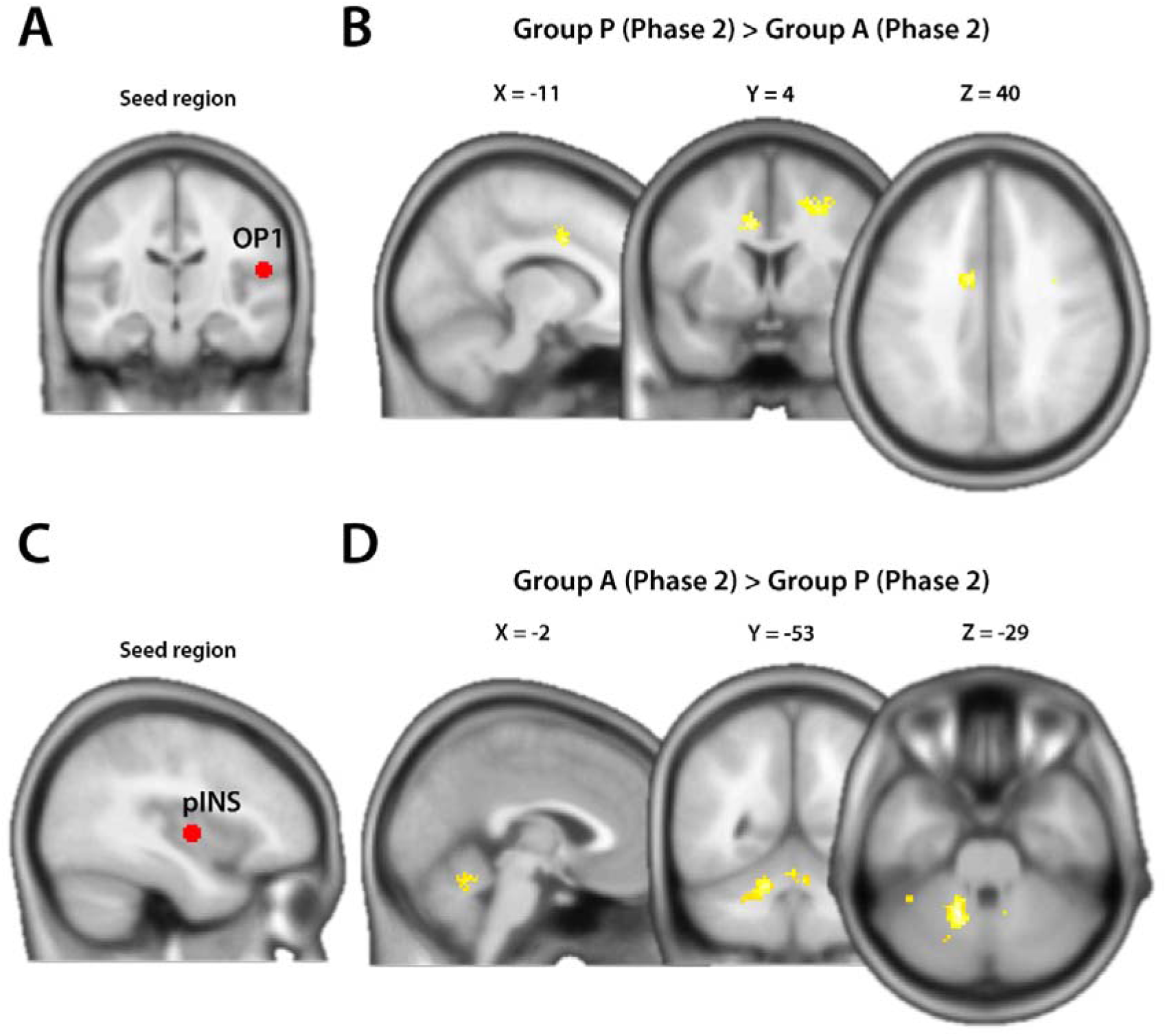
Whole-brain functional connectivity results using generalized psychophysical interactions (gPPI). The OP1 seed region (A) demonstrated an enhanced functional coupling to the aMCC during pain (B). The pINS seed region (C) showed an enhanced interaction with the cerebellar culmen during pain relief (D). q(FDR)<0.05, cluster-based corrected.

**Table 2.**
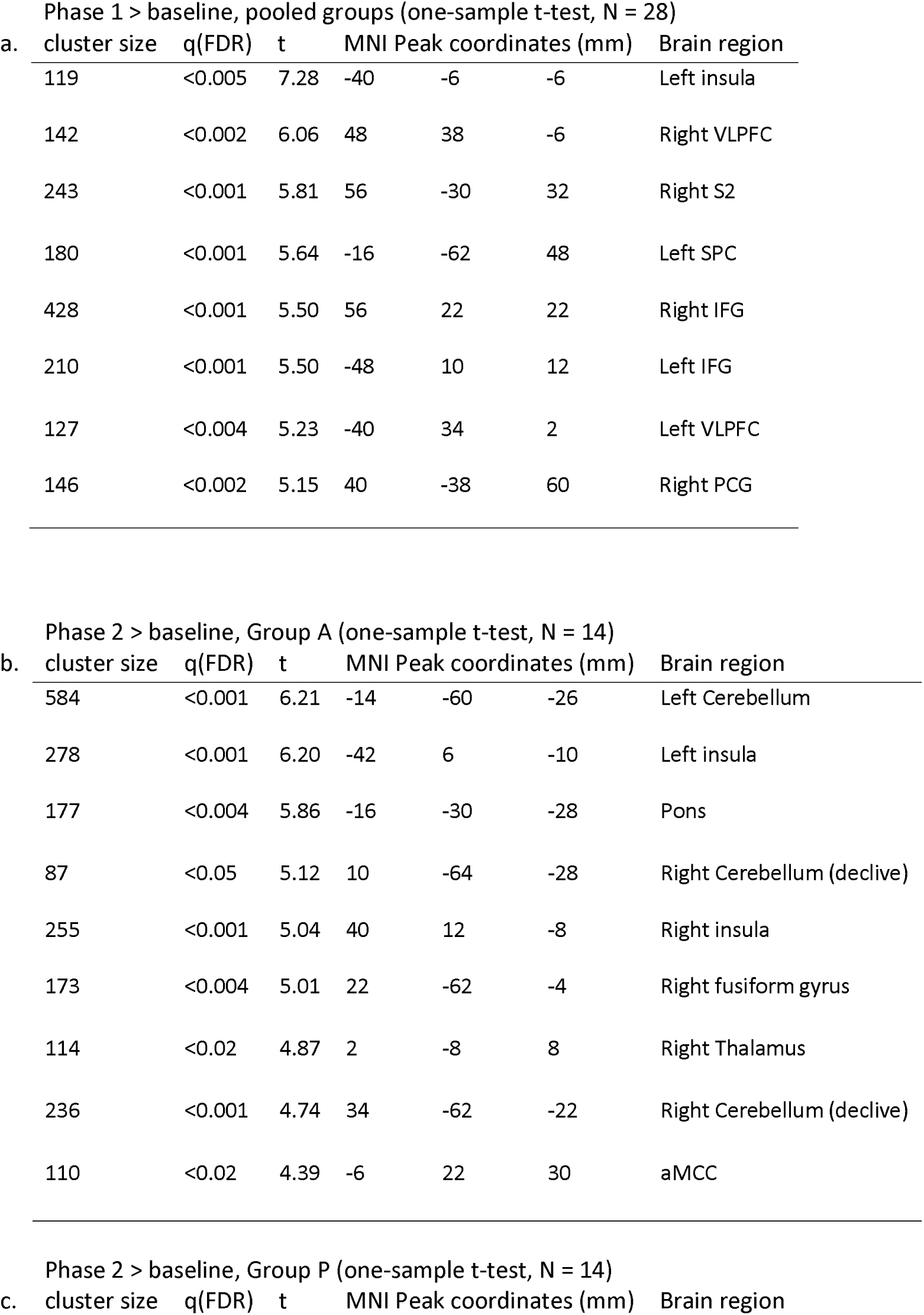
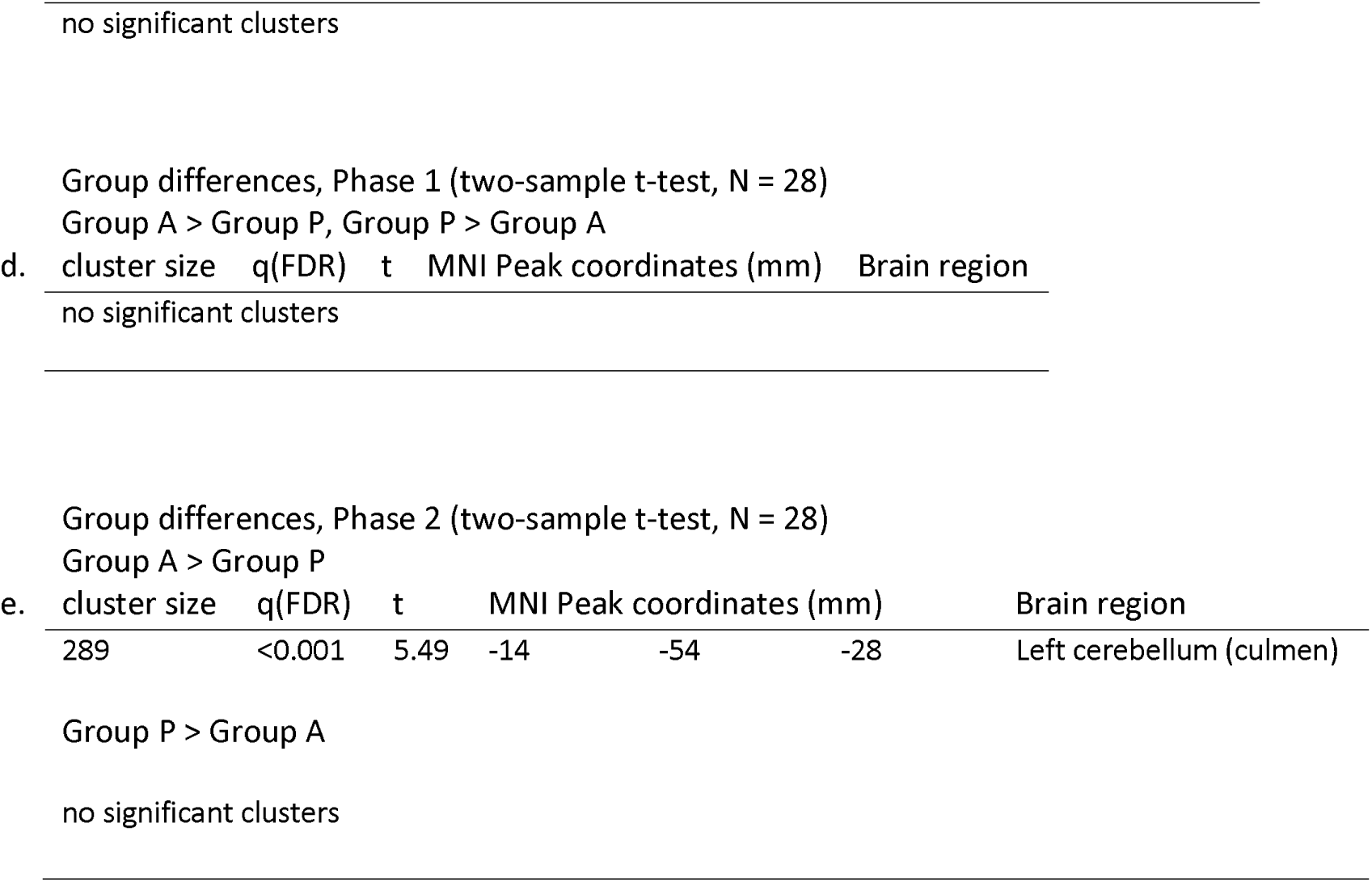
Functional connectivity results of Phase 1 and Phase 2 with pINS seed region. MNI = Montreal Neurological Institute, FDR = False discovery rate, VLPFC = ventrolateral prefrontal cortex, SPC = superior parietal cortex, S2 = secondary somatosensory cortex, IFG = inferior frontal gyrus, PCG = postcentral gyrus

In contrast, the OP1 region (Figure 3a) revealed a positive functional coupling to the bilateral S2, right supplementary motor area (SMA), bilateral inferior parietal cortex (IPC), bilateral thalamus, left posterior midcingulate cortex (pMCC), caudate, cerebellum, and right occipital cortex (OC) (q(FDR) < 0.05, Table 3a).

**Table 3.**
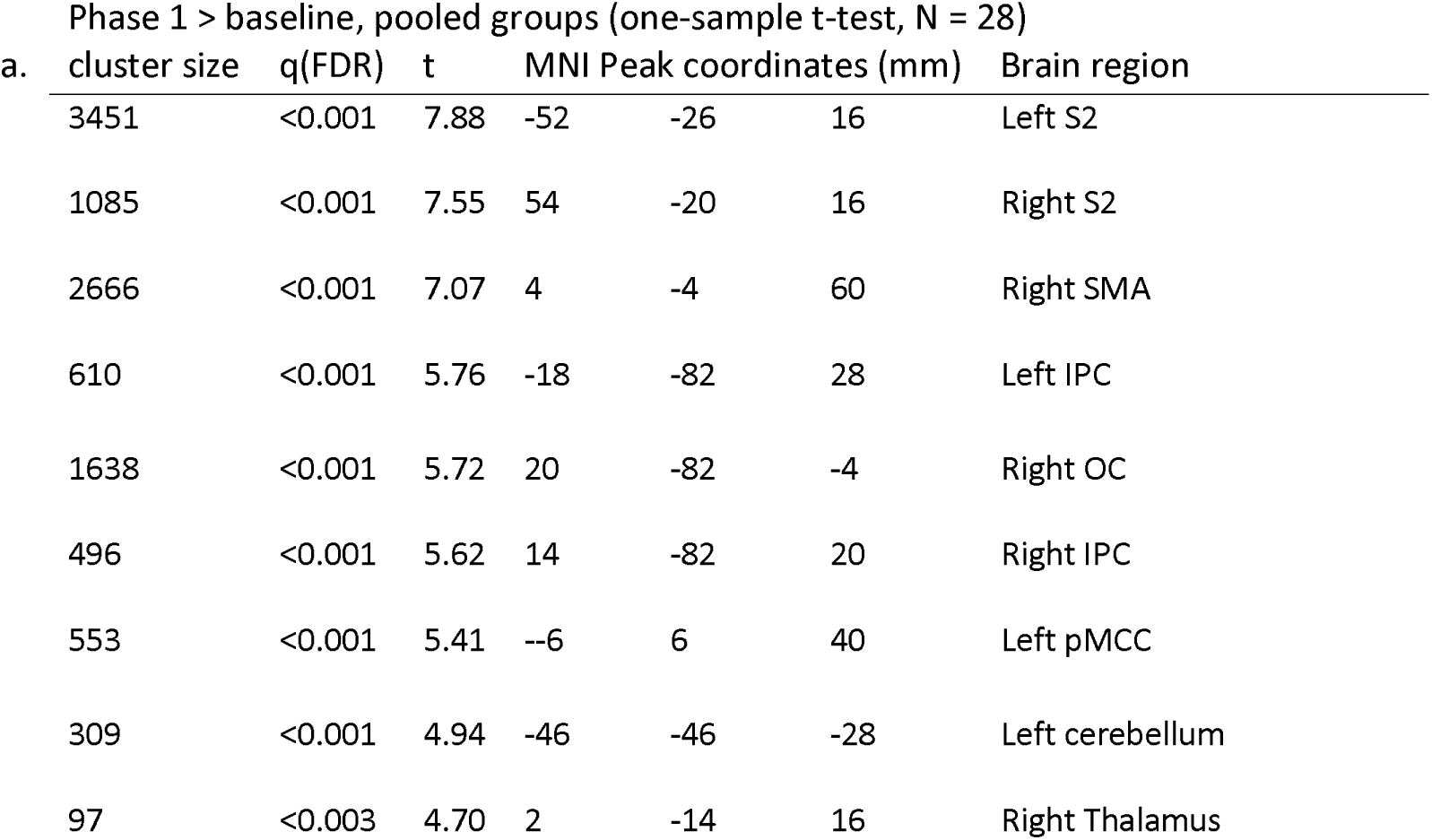
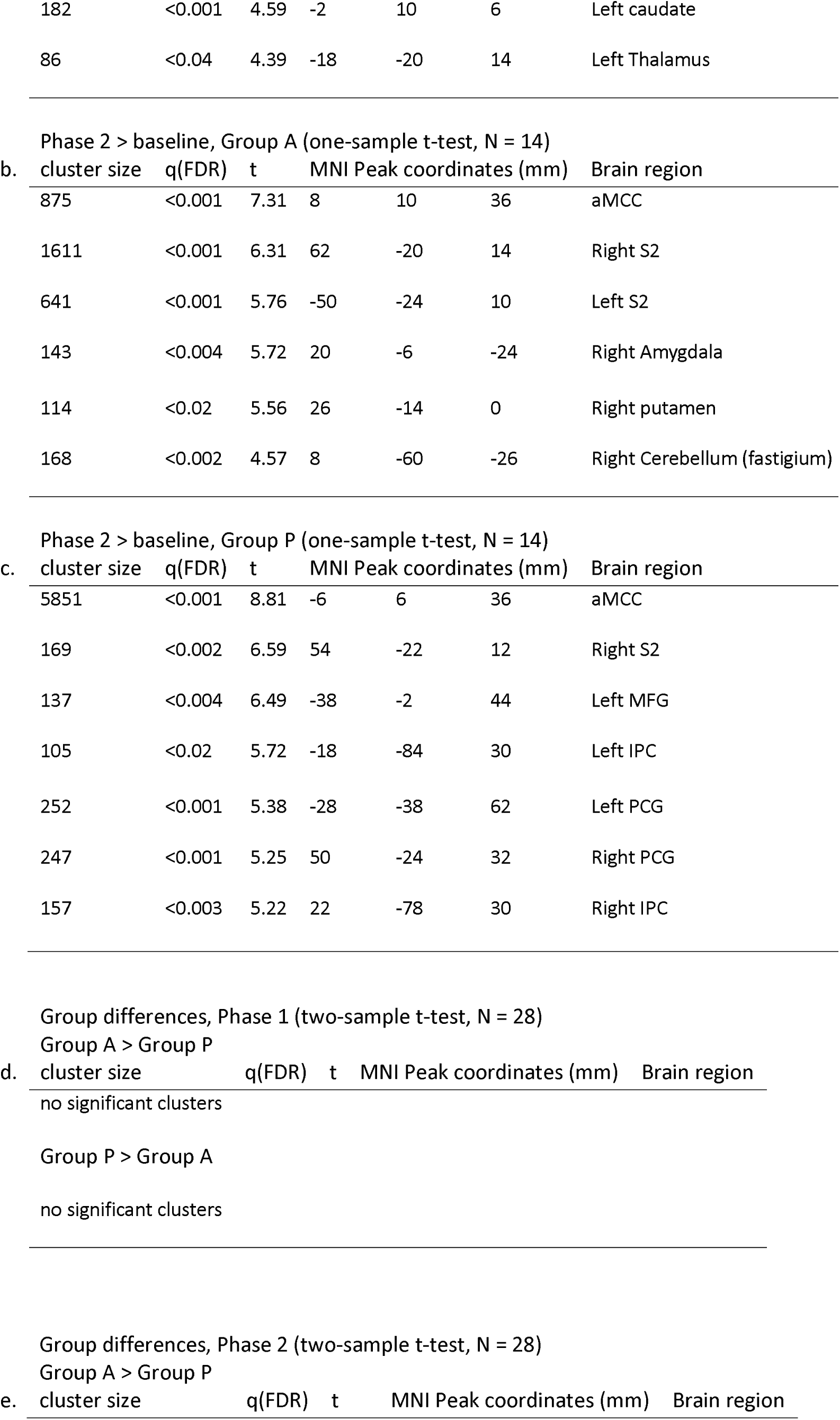
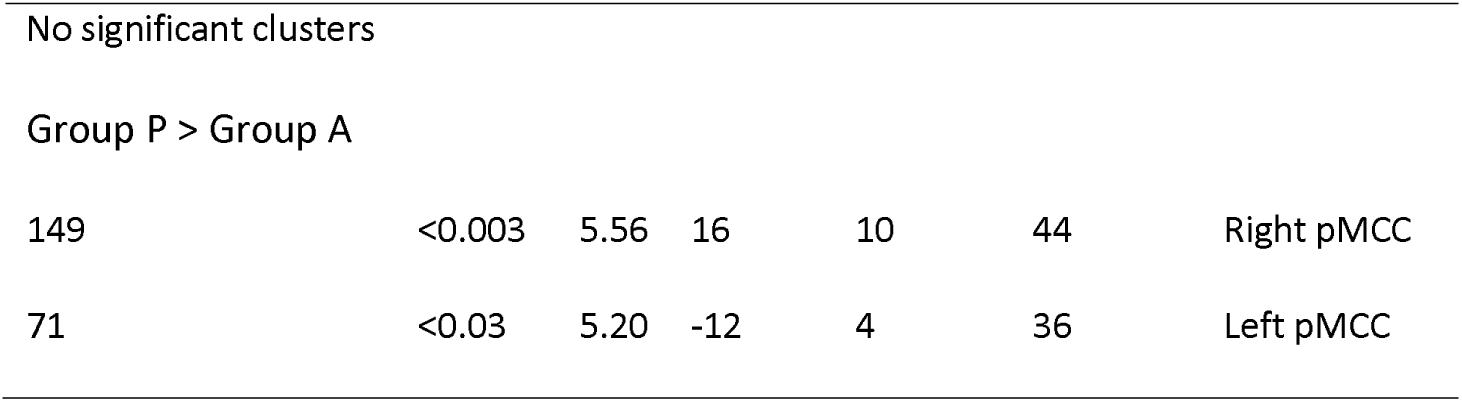
Functional connectivity results of Phase 1 and Phase 2 with OP1 seed region. MNI = Montreal Neurological Institute, FDR = False discovery rate, aMCC = anterior midcingulate cortex, pMCC = posterior midcingulate cortex, DLPFC = dorsolateral prefrontal cortex, IPC = inferior parietal cortex, S2 = secondary somatosensory cortex, SMA = supplementary motor area, OC = occipital cortex, MFG = middle frontal gyrus, PCG = postcentral gyrus

For the functional connectivity results of phase 2 within each group please see Tables 2b, 2c (for pINS) and 3b, 3c (for OP1).

#### 3.3.2 Group differences

Using the pINS as a seed, phase 1 revealed no significant group differences (Table 2d). The between-group analysis of phase 2 demonstrated a positive functional coupling of the pINS exclusively to the culmen of the left cerebellum during pain relief (q(FDR) < 0.001, Figure 3d, Table 2e). The reverse contrast, Group P (phase 2) > Group A (phase 2), revealed no significant results.

In contrast, the OP1 between-group analysis yielded a significant enhanced functional coupling to bilateral clusters of the pMCC region during noxious stimulation (q(FDR) < 0.03, Figure 3d, Table 3e). The reverse contrast, Group A (phase 2) > Group P (phase 2), revealed no significant results.

## 4. Discussion

Our study results contribute to the ongoing validation of distinct pain-related brain mechanisms (Mano and Seymour 2015; Kucyi and Davis 2015; Davis et al. 2015; Garcia-Larrea 2012b). The main finding confirms the prominent role of the operculo-insular cortex as an important node in pain processing. Furthermore, we identified novel and distinct neural interactions between the pINS/pMCC and OP1/cerebellum, indicating two functionally separate mechanisms during pain perception and relief.

### 4.1 Stimulus

The idea behind selecting a tooth as a target site for a purely nociceptive stimulus is not new (Chatrian et al. 1975) and relies on the observation that repetitive electrical stimuli reliably evoke painful short and sharp sensations (A-delta fiber-mediated pain) and no superimposed mechano- or thermosensations (Narhi et al. 1992; Brugger et al. 2011; Brugger et al. 2012; Meier et al. 2015; Meier et al. 2014). Nociceptive fibers are generally more susceptible to local analgesics than thickly myelinated fibers, as clinically demonstrated by a pressure sensation during dental extractions despite complete analgesia (Becker and Reed 2012). Correspondingly, the subjects in our experiment reported that electrical stimuli evoked a distinct non-painful sensation at the target tooth after the onset of analgesia, which permitted a task-based activity and connectivity analysis. Furthermore, adaptive changes in the perception of the painful repetitive electrical stimulus (sensitization/habituation) within the time frame of the experiment were minimal as demonstrated in our previous experiments (Meier et al. 2015; Meier et al. 2014; Brugger et al. 2012). This allowed the exclusion of a continuous pain rating task that prevented the possible blurring effects associated with cognitive and/or motor aspects of pain intensity rating (Baliki et al. 2009; Oertel et al. 2012).

### 4.2 Posterior insula and parietal operculum

The current results confirm accumulating evidence from human reports demonstrating that the operculo-insular region is the most consistently activated area in acute pain (Garcia-Larrea 2012a; Oertel et al. 2012; Duerden and Albanese 2013; Favilla et al. 2014; Segerdahl et al. 2015). In support of this observation, the pINS and the medial parietal operculum are the only areas where electrical stimulation triggered somatic pain in pre-surgical patients (Mazzola et al. 2012b). Furthermore, electrophysiological recordings revealed that the earliest brain responses to noxious stimuli originate in the operculo-insular cortex (Garcia-Larrea et al. 2003). On the other hand, by using intracerebral recordings, it has been recently shown that the pINS also responds to stimuli unrelated to nociception, which confutes the widespread assumption that the pINS serves as a pain-specific center (Liberati et al. 2016). Nevertheless, one should be cautious to not overintepret these results as a specific involvement of the pINS in pain perception cannot be excluded. In particular, as the local field potentials used in that study reflect the firing of cell populations, pain-specific neurons in the pINS cannot be excluded and further research in this area is needed. Surprisingly, the current investigation revealed an ipsilateral (left-sided) activation of the pINS. Yet, in contrast to the mainly contralateral spinothalamic input from spinal nerves, the trigeminal cortical representation is known to respond to ipsi- and bilateral receptive fields (Cusick et al. 1986; Lin et al. 1993; Jantsch et al. 2005). This is in keeping with observations from Rasmussen and Penfield who elicited ipsi- and bilateral sensations from cortical stimulation of the face and oral cavity areas in awake humans (PENFIELD 1947; RASMUSSEN and PENFIELD 1947). Supporting this observation, a minority of mandibular branch proprioceptive afferents cross the midline, and thus ascend homolaterally to the thalamus (Chen et al. 1997), and the insular region is known to receive bilateral input from the trigeminus (Dong et al. 1989). Furthermore, our results are in agreement with an experimental tooth pain study which demonstrated bilateral S2 activation and a clear dominance of tooth over hand representation in the ipsilateral insula, although the two different stimulation modalities might have contributed to this difference (tooth: electrical vs. hand: mechanical) (Jantsch et al. 2005). Nevertheless, as we only stimulated and anesthetized a single tooth, a final conclusion about lateralization effects cannot be drawn, which is an important study limitation.

Furthermore, our results indicate a pain-related pINS activation that is located more anterior to the dorsal pINS activation reported in several pain studies (Henderson et al. 2011; Garcia-Larrea 2012a; Wager et al. 2013; Segerdahl et al. 2015). However, electrical stimulations in pre-surgical patients elicited somatic pain not only in the dorsal pINS regions but also in the widely distributed anterior portions of the pINS (Mazzola et al. 2012a). Moreover, a somatotopic organization of pain within the insular cortex was suggested, with the face being represented anterior to the upper and lower limbs (Mazzola et al. 2009). In keeping with this observation, another study applying painful trigeminal CO2 stimuli to the nasal mucosa revealed a pain-related activation in a more anterior region of the pINS that is comparable to the pINS activation of our study (Oertel et al. 2012).

Regarding the human parietal operculum, it is classified into four distinct cytoarchitectonic areas (OP1-4) which can be interpreted as anatomical correlates of the functionally defined human S2 region (Eickhoff et al. 2006a; Eickhoff et al. 2006b). Our results indicate a contralateral (right-sided) neural response of the OP1 area. This is in line with a previous report that demonstrated a right-sided dominance of activation in the S2/subcentral area during bilateral painful electrical tooth stimulation (Brugger et al. 2011). A meta-analysis indicated that painful stimuli are exclusively processed in the OP1 region, whereas non-painful stimuli are represented in the OP 2-4 region (Eickhoff et al. 2006a). Further support for this observation comes from lesion studies (Greenspan et al. 1999; Greenspan and Winfield 1992). In contrast, another investigation detected pain-related effects exclusively in the OP 4 region, which raises doubts about a definitive answer regarding pain processing within the parietal operculum (Mazzola et al. 2012b).

### 4.3 Cingulate cortex

Interestingly, the OP1 region demonstrated an enhanced functional coupling with the pMCC during noxious stimulation (Figure 3D). An anatomical classification of the cingulate cortex subregions has been proposed by Vogt and colleagues (Vogt et al. 2003; Vogt 2016). Of these, the MCC plays a key role in pain processing and is almost as consistently activated as the operculo-insular cortex (Vogt 2005; Frot et al. 2008). The observed neural interaction of the OP1 region with the pMCC closely corresponds with a study using intracerebral recordings of laser-evoked potentials showing the simultaneous processing of nociceptive information in both brain regions (Frot et al. 2008). Although our results indicate a highly pain-related relationship between the OP1 and pMCC, no pain-only area has been identified in the cingulate cortex to date (Vogt 2016). Alternatively, a multidimensional view including behavioral goals such as avoiding noxious stimuli has been proposed (Vogt 2005). It is assumed that the pMCC is involved in the withdrawal reactions from the painful stimulus in maintaining connections to the motor cortex and SMA for the preparation of voluntary movements (Buchel et al. 2002; Niddam et al. 2005; Vogt 2016). Withdrawal from the pain stimulus has to be operational at very short latencies and is, therefore, not dependent on prior sequential processing through other cortical areas (Frot et al. 2008).

### 4.4 Cerebellum

Very few studies to date have attempted to unveil the cerebellum’s function in pain. A review by Moulton and colleagues summarizing animal and human reports indicated that the cerebellum receives primary nociceptive afferents and that the electrical and pharmacological stimulation of the cerebellum can modulate pain processing (Moulton et al. 2010). Further, cerebellar lesions can lead to altered pain perception (Ruscheweyh et al. 2014). Although cerebellar activity occurs consistently in the presence of acute pain (Apkarian et al. 2005; Peyron et al. 2000), little is known about the specific role of the cerebellum regarding pain perception. Recently, the posterior cerebellum has been shown to process the overlapping and multimodal inputs from motor control and pain (Coombes and Misra 2016). Animal electrophysiological studies demonstrate direct evidence of the afferent input from nociceptors to the cerebellum (VanGilder 1975; Garwicz et al. 1998; Moulton et al. 2010), e.g., A-delta fiber stimulation leads to activation in Purkinje cells in the anterior lobe of the cerebellum in cats (Ekerot et al. 1987). Furthermore, accumulating evidence from human reports demonstrates that the cerebellar processing may be more directly related to pain and nociceptive modulation (Bingel et al. 2002; Helmchen et al. 2003; Moulton et al. 2011). In support of this, noxious heat produced cerebellar activation even under general anesthesia using propofol (Hofbauer et al. 2004). By considering the current results, the noxious stimulation in phase 1 led to cerebellar activation that was widely distributed in the posterior and anterior lobes. In contrast, the pINS seed region demonstrated an exclusive functional coupling with the cerebellar culmen in the anterior lobe during pain relief, suggesting a distinct nociceptive modulatory relationship between these two brain regions. Evidence supporting this observation comes from a rat study where microinjections of morphine into the cerebellar culmen resulted in acute analgesia (Dey and Ray 1982). Moreover, a recent study identified three functional cerebellar clusters during noxious stimulation, among which one significant cluster mainly involved the culmen that was functionally connected to the bilateral posterior portions of the insula (Diano et al. 2016). These results indicate a potential role of the pINS and cerebellar culmen in basic nociceptive processing and modulation.

### 4.5 Conclusion

The neural block of trigeminal nociceptive primary afferents leads to a significant activity reduction in the pINS and OP1 regions, but not in other pain-associated brain areas. The current findings thus strengthen the evidence for the unique relevance of the operculo-insular cortex in pain perception. The pINS and OP1 seed regions seem to maintain separate neural cross-talks that might be related to pain relief (cerebellar culmen) and immediate withdrawal behavior (pMCC). However, it must be noted that these conclusions are based on reverse inference. Thus, the likelihood of the reverse inference being true is a function of the degree to which the brain mechanism (e.g. pINS – cerebellum connectivity) is exclusively triggered by the proposed psychological state (pain relief). Nonetheless, the current results support the conceptual framework that during the pain experience, localized nociceptive nodes interact with distributed brain targets.

## 5. Acknowledgments

This work was supported by a grant from the Swiss Dental Association and partly by Glaxo Smith Kline, Consumer Healthcare, Weybridge, UK. In addition, we would like to thank all volunteers who participated in the current study.

